# Transcriptomics analysis reveals intracellular mutual regulation of coral-zooxanthella holobionts

**DOI:** 10.1101/2021.11.03.467195

**Authors:** Yunchi Zhu, Xin Liao, Tingyu Han, J.-Y. Chen, Chunpeng He, Zuhong Lu

## Abstract

Corals should make excellent models for cross-kingdom regulation research because of their natural animal-photobiont holobiont composition, yet a lack of studies and experimental data restricts their use. Here we integrate new full-length transcriptomes and small RNAs of four common reef-building corals with the published *Symbiodinium C1* genome to gain deeper insight into mutual gene regulation in coral-zooxanthella holobionts. We show that zooxanthellae secrete miRNA to downregulate rejection from host coral cells, and that a potential correlation exists between miRNA diversity and physiological activity. Convergence of these holobionts’ biological functions in different species is also revealed, which implies the low gene impact on bottom ecological niche organisms. This work provides evidence for the early origin of cross-kingdom regulation as a mechanism of self-defense autotrophs can use against heterotrophs, sheds more light on coral-zooxanthella holobionts, and contributes valuable data for further coral research.

## Introduction

miRNA is a type of small, single-stranded non-coding RNA found in plants, animals, and some viruses, and that functions in RNA silencing and post-transcriptional regulation of gene expression (1). miRNA is recognized to be involved in various regulation pathways, such as cross-cell regulation, cross-organ regulation, and even cross-generation regulation, however, whether cross-kingdom regulation occurs has been controversial. In 2012, Zhang and colleagues first proposed this concept after they identified stable plant miRNAs in several human and animal organs (2). They argued that intaken plant miRNAs could remain active and exert biological effects in animal tissues. A great deal of subsequent research has supported their views, for example, (3) revealed a correlation between fruit juice intake and fruit miRNA concentration in human serum; one honeysuckle miRNA (4) and two rapeseed miRNAs (5) were observed to be effectively absorbed by mice; (6) integrated ddPCR and transcriptome analyses to explore silkworms’ absorption of mulberry miRNAs; etc. Some “avid” supporters of the idea of cross-kingdom miRNA activity advanced the medicinal value of plant miRNA against IAV (4), tumors (7), and even SARS-CoV-2 (8). Nevertheless, ever since the concept was put forward, the doubts continue to be voiced (9, 10). The proposers and supporters of cross-kingdom regulation have been working on explaining miRNA absorption mechanisms in detail in order to refute these objections (11), while not all research of interorganismal miRNA exchange is focused on food-intake-derived regulation, such as (12) studied the role of dodder miRNA in the parasitic process.

Most studies on cross-kingdom miRNA activity study humans or other complex animals as experimental subjects and look into whether plant miRNAs can affect animals’ physiological functions. From our perspective, these models may be too complex in terms of behavior and physiology to support accurate conclusions about the origins or even the existence of such small molecules. Instead, we suggest a focus on more basal species exhibiting less complex behavior to explore the origins and details of the dominant regulation mechanisms of miRNA, which could lay the theoretical and practical foundation for studies of more complex organisms. We recommend coral as a potential animal model for cross-kingdom regulation research. Known co-evolutionary patterns exist for coral microbial communities and coral phylogeny. Corals obtain the majority of their energy and nutrients from photosynthetic unicellular dinoflagellates of the genus *Symbiodinium*, commonly known as symbiotic zooxanthellae. Zooxanthellae living within corals’ tissues not only supply their hosts with the products of photosynthesis but also aid in coral calcification and waste removal. Coral-zooxanthella holobionts are naturally formed “animal-plant mixtures,” that offer the advantages of simple behavior and ease of observation relative to more complex animal or plant models. It is not difficult to assume that cross-kingdom regulation, regarded by its proposers as evidence of long-term co-evolution between photosynthetic organisms and animals, may play an important role in these holobionts.

However, in contrast to more common model organisms, there is a lack of public omics data from reef-building corals, not commensurate with their important status in the marine ecosystem and scientists’ sharp focus on them. At the end of 2020, only 24 coral genomes at a medium assembly level are available on NCBI, while transcriptomes and proteomes are even more lacking. Many other well-known databases, such as miRBase (13) and KEGG (14), contain very little or even no coral data. Various factors, including geographic location, technical limitations, and experimental challenges, contribute to the limited number of coral studies serving as data sources, resulting in a “data gap.” This lack of data deprives bioinformaticians of the opportunity to conduct dry lab work on corals, adding to a vicious circle exerting further negative effects on coral research. In-depth research underpinned by high quality bio-data is in urgent demand to break this cycle.

In order to explore intracellular mutual regulation of coral-zooxanthella holobionts, we sequenced the full-length transcriptomes and small RNAs of four common and frequently dominant reef-building corals, including *Acropora muricata, Montipora foliosa, Montipora capricornis* and *Pocillopora verrucosa*. For each species, we also performed Illumina sequencing to quantify gene expression. Every full-length transcript was annotated with Nr, Nt, Pfam, KOG, Swiss-Prot, GO and KEGG to distinguish coral and zooxanthella genes. A public *Symbiodinium C1* genome was employed to identify zooxanthella miRNAs, whose targeting sites were predicted later. Finally, highly expressed coral genes, zooxanthella genes and miRNA-target genes were selected for enrichment analysis. To a certain extent, zooxanthellae become organelles when they are inside coral cells, limiting the expression of most of their genes. Some of the miRNAs secreted by zooxanthellae tend to target immune-related genes in corals, suggesting that cross-kingdom regulation be a way for the zooxanthellae to prevent rejection by their hosts. Results also demonstrate that coral genes turn out to be functionally similar despite structural differences among coral species. These results suggest that the experimental subject selection for a coral model organism may be flexible, undoubtedly a benefit for researchers. This work sheds light into the coral-zooxanthellae holobiont, an ancient association in which an early example of cross-kingdom regulation is found. We also curate our data in a full-service online database to support further research (http://coral.bmeonline.cn).

## Results

All raw sequencing data are available on NCBI (**Table 1**). 28 transcriptomes have been sequenced in total. For each species, the biological replicate numbers of full-length SMRT transcriptome sequencing, small RNA sequencing and Illumina RNA-seq sequencing are designated as 1:3:3 in order to ensure accuracy.

**Table 1.**
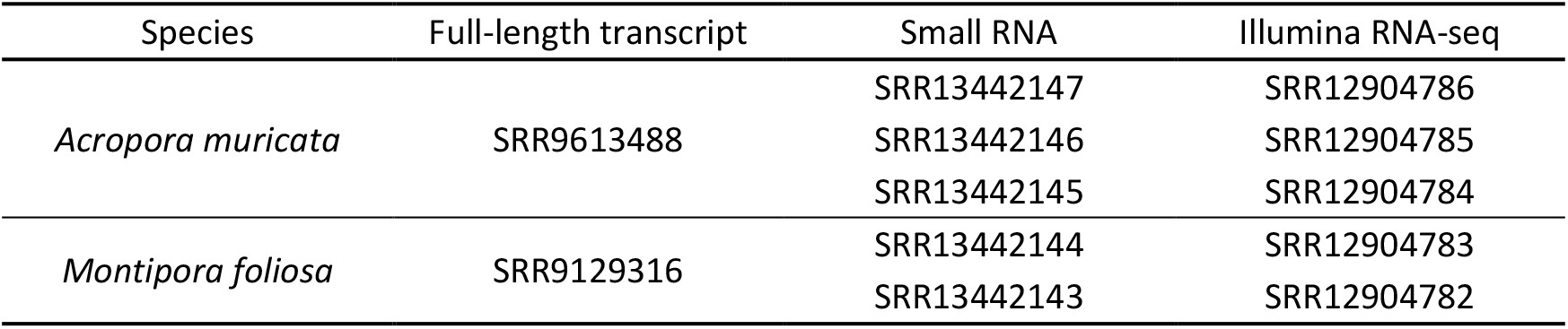
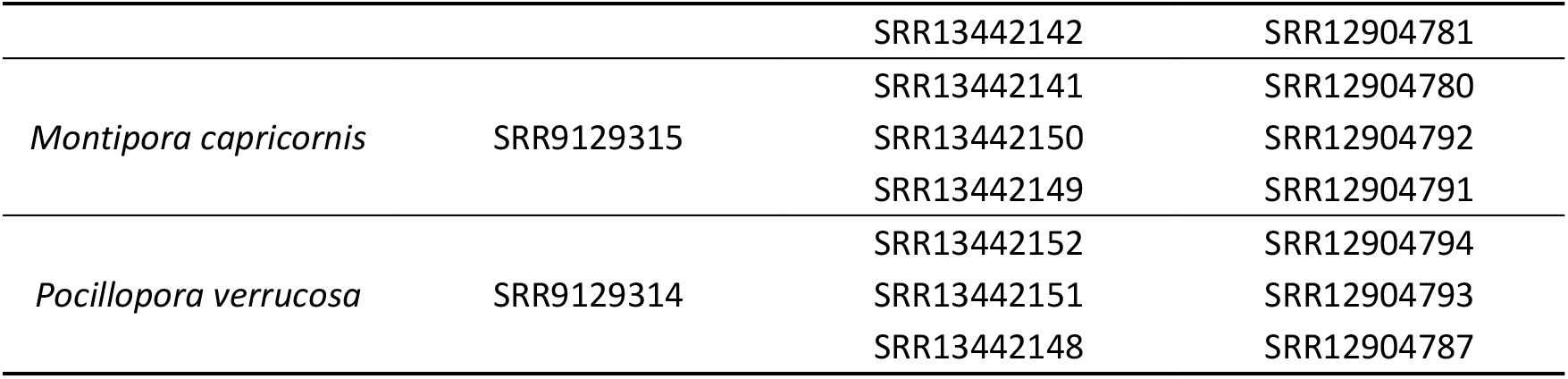
Raw sequencing data on NCBI SRA. There are 28 transcriptomes in total. Each sample represents one biological replicate.

### Transcriptome sequencing and data processing

The SMRT-sequencing technology was performed with the PacBio Sequel II platform to acquire offline polymer read bases of full-length transcriptomes using SMRTlink v. 7.0 software. The offline polymer read bases of *A. muricata, M. foliosa, M. capricornis* and *P. verrucosa* samples are 12.6G, 20.16G, 16G and 14.97G respectively (**File S1** for more details). The subreads, CCSs, FLNCs, consensus sequences, corrected consensus reads and unigenes are shown in **Table 2**, which also covers all information revealed in subsequent analyses.

**Table 2.** Sequencing data statistics of four full-length coral transcriptomes.

| Sample name | *A. muricata* | *M. foliosa* | *M. capricornis* | *P. verrucosa* |
| --- | --- | --- | --- | --- |
| Subread base (G) | 12.37 | 19.82 | 15.62 | 14.61 |
| Subread (number) | 5,034,975 | 7,847,499 | 7,535,422 | 7,357,796 |
| Average subread length (Nt) | 2,456 | 2,526 | 2,073 | 1,986 |
| CCS (number) | 358,971 | 403,628 | 354,221 | 317,878 |
| FLNC (number) | 264,098 | 291,254 | 248,892 | 236,923 |
| Consensus read (number) | 22,925 | 23,067 | 18,354 | 20,696 |
| Corrected consensus (number) | 22,925 | 23,067 | 18,354 | 20,696 |
| Unigene (number) | 14,926 | 14,040 | 11,410 | 12,982 |

The full-length transcriptomes were annotated with the aid of Nr, Nt, Pfam, KOG, Swiss-Prot, GO and KEGG databases, and related unigene statistics are shown in **Fig 1**. In function-related databases, over 70% of unigenes get GO and KEGG annotations, paving the way for functional enrichment analysis. In protein-related databases, more than 96% of the unigenes of investigated corals are annotated in Nr, the basic protein primary sequence database. Such wide coverage indicates that Nr could be utilized for gene-ID mapping among different corals. As illustrated in **Fig S1**, Nr annotations map most genes of the four corals to the cnidarians *A. digitifera, E. pallida* and *N. vectensis*, with more than 85% overlap. Acceptable as it is, this result still reflects the lack of coral data for those annotations rarely correspond with the actual species in question.

**Fig 1.**
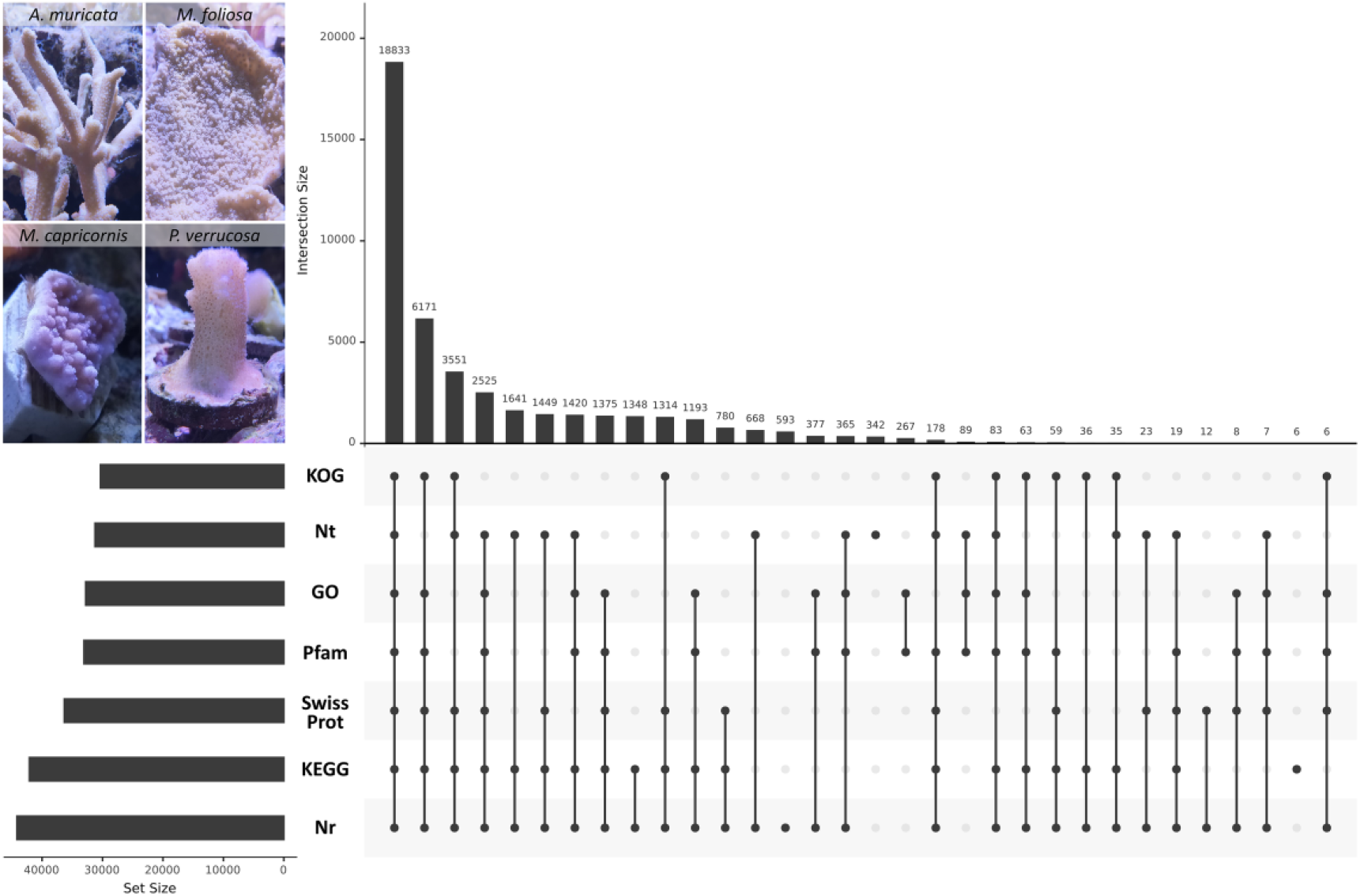
Statistics of gene functional annotation. Photos of investigated corals are listed in the upper left corner.

According to Nr annotation, there are 43 *Symbiodinium* genes in *A. muricata*, 159 in *M. foliosa*, 45 in *M. capricornis*, and 129 in *P. verrucosa*, less than 0.07% of the total gene number. These numbers should be described as significantly few, although errors in annotation databases could account for this phenomenon to some extent. This result is similar to the condition of several well-researched organelles, such as mitochondria and chloroplasts, that are hypothesized to have become “intracellular slaves” during the “war” between archaebacteria and proteobacteria one billion years ago, working all the time but hardly expressing any genes of their own. It is not unreasonable to wonder whether zooxanthellae have undergone an analogous situation in the holobionts.

The RNA-seq sequencing was performed on the Illumina HiSeq X Ten platform. The total read bases are 22.1G (7.8G + 7.2G + 7.1G) from *A. muricata*, 29G (9.9G + 8.4G + 10.7G) from *M. foliosa*, 23.2G (7G + 7.1G + 9.1G) from *M. capricornis* and 23.1G (6.9G + 7.5G + 8.7G) from *P. verrucosa*.

Small RNAs were sequenced with the Illumina HiSeq X Ten platform as well. The raw read bases are 4.412G (1.548G + 1.359G + 1.505G) from *A. muricata*, 3.865G (1.275G + 1.053G + 1.537G) from *M. foliosa*, 3.295G (1.177G + 0.893G + 1.225G) from *M. capricornis* and 6.235G (2.67G + 1.494G + 2.071G) from *P. verrucosa* (**File S2** for more details). The raw read, clean read, unique sRNA, rRNA read, tRNA read, snRNA read, snoRNA read, known coral miRNA, novel coral miRNA and novel Symbiodinium miRNA are shown in **Table 3**.

**Table 3.**
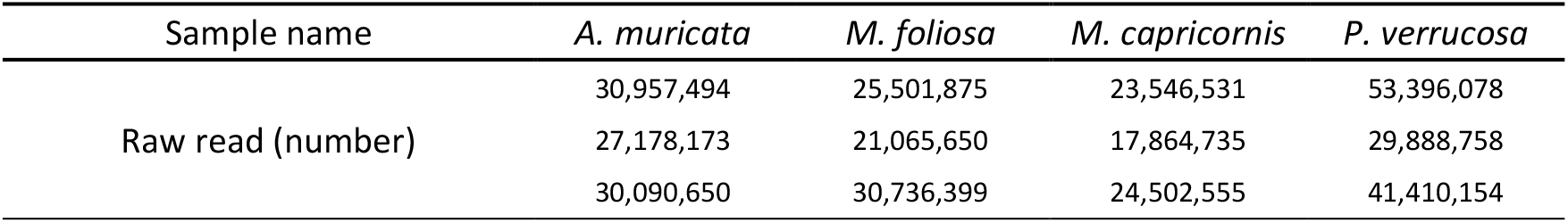
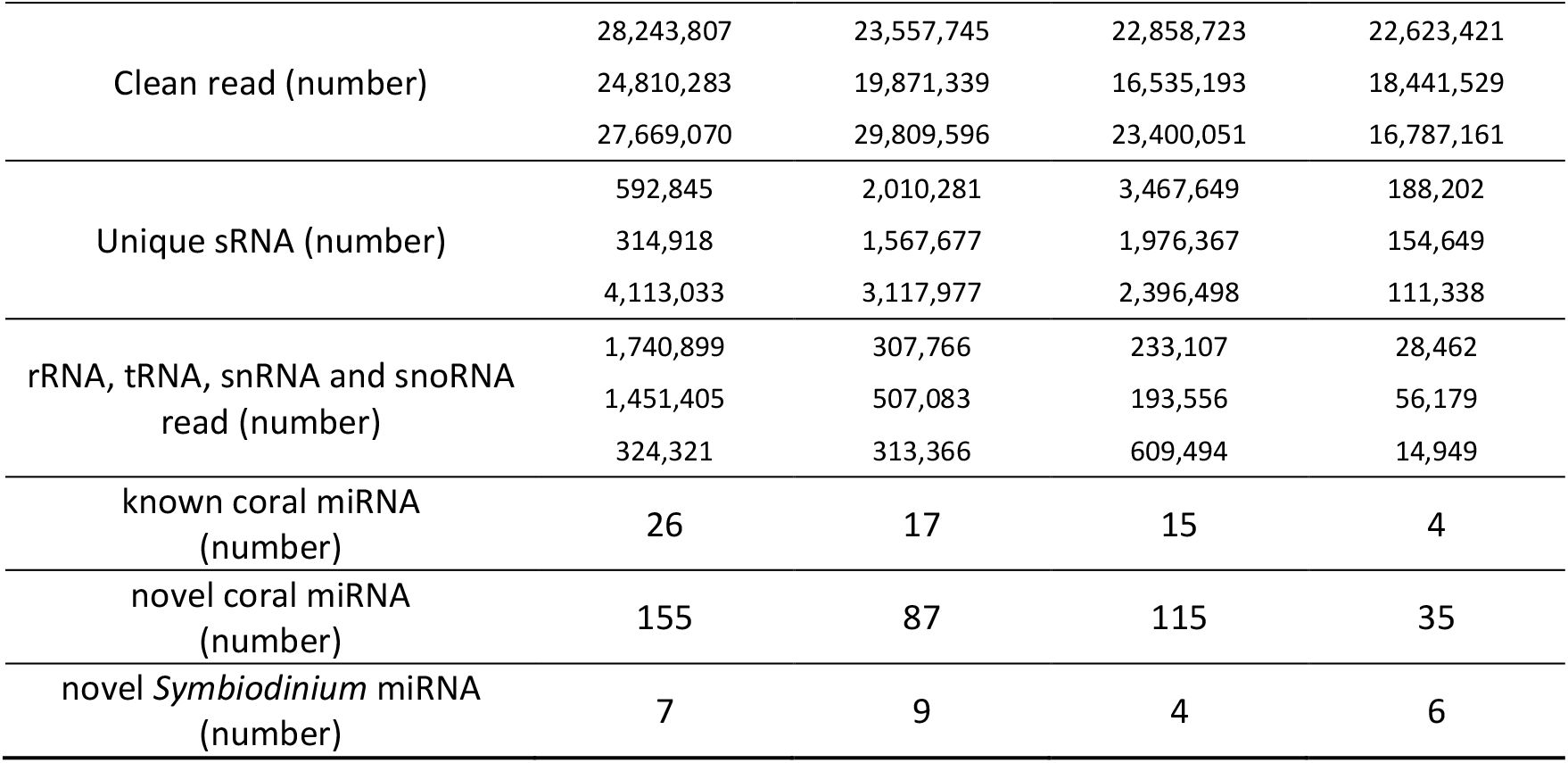
Sequencing data statistics of four coral small RNAs. For each species, miRNA identification results of three biological replicates were combined.

436 coral miRNAs and 11 novel *Symbiodinium* miRNAs are identified, the detailed information of which, such as loci, sequence and precursor structure, have been added to corresponding miRNA datasets named CoralmiR (http://coral.bmeonline.cn/miR/) and CoralSym (http://coral.bmeonline.cn/sym/). Comparing the two, CoralmiR is large yet flawed in that there are inevitably several false positive results in it due to the de novo analysis process, while CoralSym is small but precise because CoralSym miRNAs are supported by published zooxanthellae genomes, enhancing their credibility.

### Gene expression analysis

Illumina RNA-seqs were mapped to full-length transcripts using RSEM for gene expression quantification and differential expression pattern identification. The results of gene expression analysis are demonstrated in **Fig 2**. Gene numbers of differential expression patterns are illustrated in **Fig 2.a**, where patterns representing differentially expressed genes in single species occupy the top three spots, while no Pattern01 gene co-expressed among all species is identified. Combined with the gene distribution revealed in **Fig 2.b**, the unique genes account for the majority of total genes in quantity and expression for each species, suggesting that gene structural differences among different corals may be significant. **Fig S2** displays the distribution of gene expression and quantity in specific sub-sets, and these supplemental results are consistent with the above findings for total gene expression.

**Fig 2.**
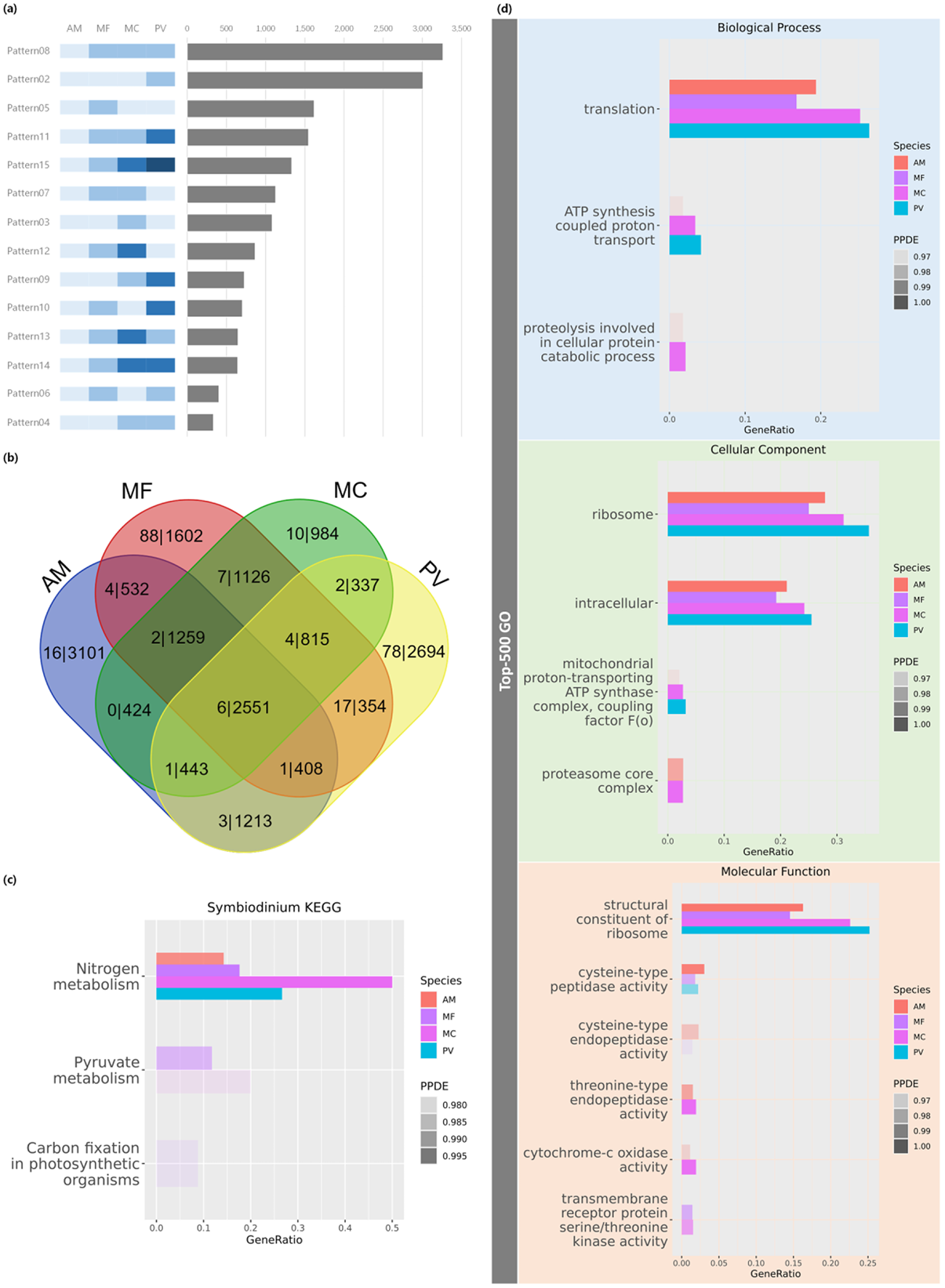
Results of gene expression analysis. AM, MF, MC, PV are short for *A. muricata, M. foliosa, M. capricornis*, and *P. verrucosa*. In the enrichment plots, each species is indicated by a different colour and statistical significance is indicated by transparency. (a) Statistics of gene differential expression patterns among the four coral species. The vertical axis is the pattern name and the horizontal axis is the number of genes. Each pattern is illustrated with a contrasting color band listed beside the vertical axis. The top three patterns in quantity are Pattern08, Pattern02 and Pattern05, which represent genes expressed differentially in *A. muricata, P. verrucosa* and *M. foliosa*, while all Pattern01 genes identified as co-expressed among species fail to meet the statistical significance criteria (PPDE>0.95). (b) Venn graph of gene distribution among four species. Text is in the form of “*zooxanthellae gene number* | *holobiont gene number*.” (c) KEGG enrichment of *Symbiodinium* genes. Pathways related to photosynthesis are put on the plot. Nitrogen metabolism is outstanding in both quantity and statistical significance, while few other pathways exhibit enrichment. (d) GO enrichment of highly expressed coral-zooxanthella holobiont genes. Significantly enriched terms are “translation” in Biological Process, “ribosome” in Cellular Component and “structural constituent of ribosome” in Molecular Function, presenting the features of housekeeping genes.

However, **Fig 2.c** and **Fig 2.d** reveal gene functional similarities among the corals in spite of their structural differences. Among the zooxanthella genes, as **Fig 2.c** indicates, nitrogen metabolism is the common and dominant pathway. Considering the large number of pathways involving photosynthesis, zooxanthella gene functions seem fairly narrow, which is reminiscent of mitochondria, in that they obtain most of the proteins they need from their host cells rather than producing them on their own. This finding may be conducive to the hypothesis that zooxanthellae have effectively degenerated into coral organelles. Highly expressed coral genes are significantly enriched in the basic cellular functions, as shown in **Fig 2.d**. The top GO terms are “translation” in Biological Processes, “ribosome” in Cellular Components and “structural constituent of ribosome” in other words, the most highly expressed genes in the corals studied are housekeeping genes. Based on the enrichment results above, genes of coral-zooxanthella holobionts from the same habitat are believed to play analogous roles that are not significantly affected by species-specific structural differences, which may help to simplify the selection of a particular coral as an experimental model subject.

**Table 4** lists the results of a hypothetical test of zooxanthella gene expression against total gene expression. Most zooxanthella genes appear to be limited in expression, hinting at suppression by host coral cells, while those from *M. foliosa* present abnormally high levels of expression in all three biological replicates. These differences provide a potential entry point for exploring cross-kingdom regulation mechanisms in coral-zooxanthella holobionts.

**Table 4.** Hypothetical test of zooxanthella gene expression against total gene expression. T-test method was utilized. 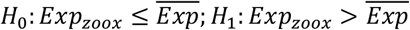

| Species | $P_{rep1}$ | $P_{rep2}$ | $P_{rep3}$ |
| --- | --- | --- | --- |
| *A. muricata* | 0.2066 | 0.0558 | 0.1367 |
| *M. foliosa* | 0.0007 | 0.0012 | 0.0014 |
| *M. capricornis* | 0.991 | 0.9934 | 0.988 |
| *P. verrucosa* | 0.3783 | 0.5881 | 0.4764 |

### miRNA-target analysis

Eleven *Symbiodinium* miRNAs are identified from the total small RNA, as shown in **Fig 3**, numbered as smg-miR-1 to 11. Their expression profiles are supplemented in **Table S2**. Four miRNAs, smg-miR-1, 2, 4, and 7, exist in all investigated corals, and their expression levels are relatively high. It is as expected that, in intracellular environments, photobiont miRNA can maintain a certain concentration far beyond concentrations observed in traditional cross-kingdom research. This phenomenon may illustrate the advantage of coral as an observer-friendly model organism to some extent.

**Fig 3.**
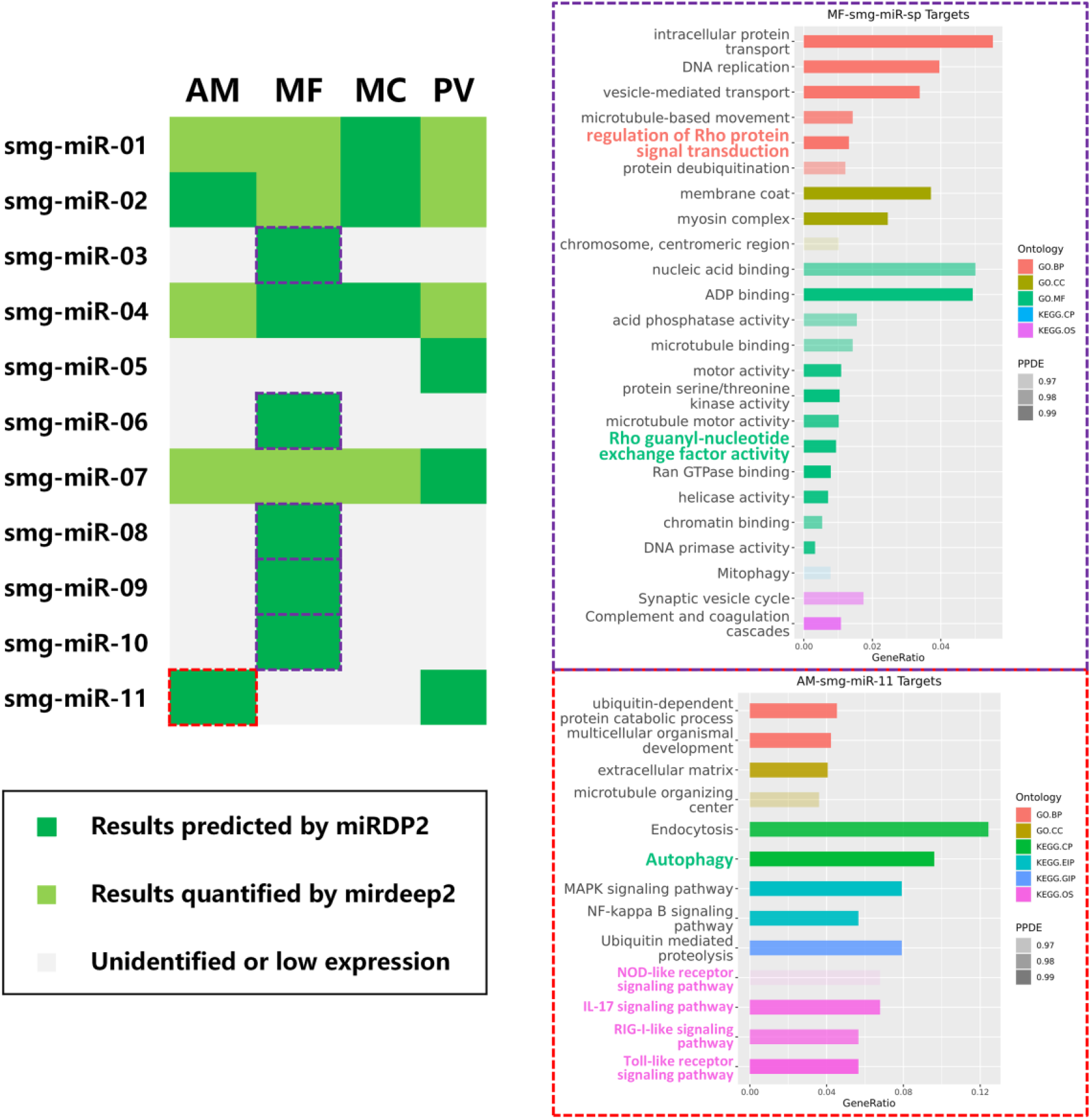
Results of miRNA-targets analysis. AM, MF, MC, PV are short for *A. muricata, M. foliosa, M. capricornis*, and *P. verrucosa*. GO.BP, GO.CC, GO.MF are short for biological process, cellular component and molecular function in gene ontology. KEGG.CP, KEGG.EIP, KEGG.GIP, KEGG.OS are short for cellular processes, environmental information processing, genetic Information processing and organismal systems in KEGG. Enrichment plots are generated with categories marked by color and statistical significance marked by transparency. Left: identified *Symbiodinium* miRNAs in investigated corals. The vertical axis indicates the miRNA name and the horizontal axis is the species name. Results of miRDP2 are marked with dark green while expanded results of mirdeep2 quantifier are marked with light green. Upper right: enrichment results of smg-miR-3,6,8,9,10 targets in *M. foliosa*. Notable terms include a series of GO terms related to cytoskeleton as well as cell migration, cell adhesion, and cell proliferation, including “regulation of Rho protein signal transduction,” “DNA replication,” and “microtubule-based movement” in biological process, “myosin complex” and “chromosome centromeric region” in cellular component and “Rho guanyl-nucleotide exchange factor activity,” “Ran GTPase binding,” “motor activity,” and “chromatin binding” in molecular function. Lower right: enrichment results of smg-miR-11 targets in *A. muricata*. Enriched pathways are immune-related pathways including NOD-like receptor signaling pathway, IL-17 signaling pathway, RIG-I-like signaling pathway and Toll-like receptor signaling pathway.

As noted in **Fig 3**, *M. foliosa* has a unique miRNA group consisting of smg-miR-3, 6, 8, 9 and 10. In the results of enrichment analysis of their target genes, a series of GO terms related to the cytoskeleton as well as to cell migration, cell adhesion, and cell proliferation emerge, among which Rho-related terms are notable. Rho GTPases belonging to the Ras superfamily of small GTP-binding protein serve as “molecular switches” in various signaling pathways, mainly by exerting effects on the cytoskeleton (15). They are widely distributed in immune-related cells and are frequently observed to participate in immune regulation. They usually cycle between an active GTP-bound state and an inactive GDP-bound state, triggering immune responses when bound to GTP until they are hydrolyzed into the GDP-bound state. Three proteins are recognized to regulate this cycle, among which Rho guanyl-nucleotide exchange factor (Rho-GEF) contributes to the activation of Rho GTPases, playing the role of positive regulator. From this it is assumed that zooxanthellae secrete these miRNAs to downregulate the activity of positive regulators in immune pathways. These miRNAs may also help remodel the cytoskeleton of corals to allow for the entry of zooxanthellae, in a similar way to how some parasites affect their hosts (16, 17).

Rho-related biological processes and molecular functions are also enriched in the target genes of smg-miR-1 in *M. capricornis* and smg-miR-7 in *P. verrucose*s shown in **Table S1**. For *A. muricata*, there is only one unique *Symbiodinium* miRNA, smg-miR-11. As illustrated in **Fig 3**, several cytoskeleton-related terms appear in GO enrichment results, while KEGG enrichment results directly demonstrate pathways in immune system. Though KEGG annotation may not be exactly terminologically accurate for corals, in that most KEGG data sources are from humans and other dominant model organisms, the results can be considered as further evidence of immune-related cross-kingdom regulation in coral-zooxanthella holobionts. In addition, autophagy-related genes are identified as these miRNAs’ targets as well, which may strengthen the credibility of the assumptions outlined above in view of many observations that Rho GTPases upregulate autophagosome fusion and transport. **Fig 4** displays a predicted pathway by which zooxanthellae block autophagosome production via miRNA-silencing the upstream Rho “switch,” nevertheless the possibility that zooxanthellae miRNAs directly target autophagy-related genes cannot yet be ruled out.

**Fig 4.**
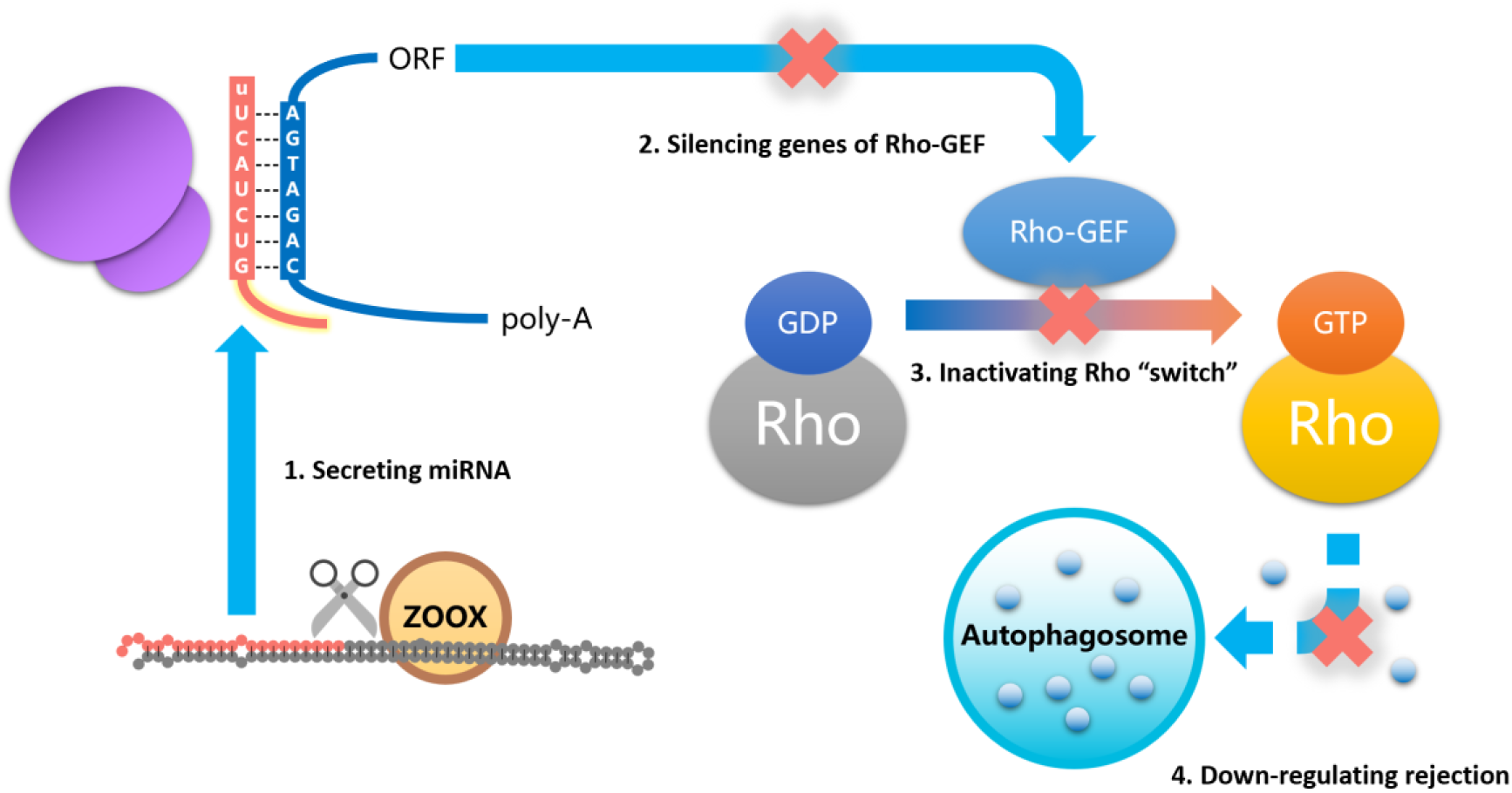
Predicted downregulating rejection pathway in coral-zooxanthella holobionts. Zooxanthellae secrete miRNAs targeting and silencing coral Rho-GEF genes to prevent the activation of Rho GTPases to block threatening autophagosome production in the coral.

In contrast to the similarities among genes in different species of corals examined in this study, zooxanthella miRNAs present functional differences among species. For instance, smg-miR-11 exists in *P. verrucosa*, while few significant functions are enriched in its target genes. smg-miR-1, 2, 4 and 7 are observed to be co-expressed among all coral samples, while their target genes are not functionally centralized. It seems that immune downregulation is the rare common role they play in the investigated corals.

No coral miRNA targeting sites are identified in the genes of zooxanthellae. It is thus inferred that miRNA silencing is not the main way corals regulate zooxanthellae, yet real matching criterion of *Symbiodinium* targeting sites may not be as strict as that applied in this research.

## Discussion

### Role of miRNAs in coral-zooxanthella holobiont

Our work provides evidence for an early origin of cross-kingdom regulation as a self-defense mechanism of autotrophs against heterotrophs. It is proposed that miRNAs target and silence immune-related coral genes to reduce rejection, helping zooxanthellae to move beyond provoking an antigenic response in potential coral hosts to becoming components of the holobiont. Conversely, no coral miRNAs are found to target zooxanthella genes, indicating that coral does not utilize miRNA as its main way to regulate zooxanthellae.

In view of the differences in gene function and expression levels revealed in **Fig 2.c** and **Table 4**, it could be further assumed that miRNA multiplicity may be conducive to function diversity for zooxanthellae. For example, among the investigated corals, the unique miRNA group in *M. foliosa*’s zooxanthellae is rather highly expressed and has diverse functions, while the miRNA in *M. capricornis*’s zooxanthellae, none of which is unique, are expressed at the lowest level. The group targeting effect of miRNA may reduce the inhibition of zooxanthellae by host coral cells so as to permit greater flexibility in the physiological activity of zooxanthellae. Records in Coral Trait (18) indicate that *M. foliosa* may be able to inhabit mesophotic coral reefs (19) more than 30m deep (https://coraltraits.org/species/1014/traits/92), where few symbiont zooxanthellae would be able to photosynthesize. The mutual dependence between coral and zooxanthellae in this holobiont under these conditions would be much lower, not only conforming to our finding but also supporting our hypothesis.

### Coral model and “ecological niche effect”

From the experimental point of view, when conducting studies of miRNA interactions between photobiont and animals, corals as model organisms have advantages in observer-friendliness and simplicity of behavior. For one thing, the single-cell structure of zooxanthellae narrows the observation range down to the intracellular environment and reduces miRNAs lost by transport and degradation, helping to maintain a relatively higher photobiont miRNA concentration for easier identification. For another, traits and behaviors of corals have been well-researched, while the internal cross-kingdom regulation occurring inside corals appears to be to be unidirectional from zooxanthellae to coral according to our research, thus it will not be too difficult to record the impact of zooxanthellae on corals under appropriate experimental conditions.

The coral-zooxanthella holobionts investigated in this study appear to have simple behavior. The majority of highly expressed coral genes are devoted to basic biological processes, such as the construction of ribosomes, while most exogenous zooxanthellae work in specific pathways under various limitations. Though our work has not identified any miRNA-based coral-to-zooxanthellae regulations, corals’ inhibitory effects on zooxanthellae are certainly observed. It is not unreasonable to predict that similar intracellular “dystopias” exist in a variety of ancient autotroph-heterotroph holobionts in which mutual regulations between symbiotic organisms vary in mechanism and are imbalanced in intensity.

It is further suggested that the closer the organism is to the bottom ecological niche, the greater the non-gene impact on it. Though coral reefs are often made up of a community exhibiting significant species diversity, our research shows that corals from the same habitat share many physiological commonalities. The “ecological niche effect” hypothesis is expected to support the feasibility of this coral model and provide guidelines for specimen collection for further research.

### Limitation and prospect

This work is typical omics-based research, mining information from large-scale bio-data, thus it is impossible to avoid systematic errors coming from bioinformatics tools and databases, especially when taking the “coral data gap” into account. Insufficient or incorrect data about coral in bio-databases affecting annotations, tags and reference sequences have emerged more than once during this research. However, the largest limitation to this research project actually comes from the lack of data about zooxanthellae. There exist various sub-clades of *Symbiodinium* in one coral, the genome differences among which may be as huge as that between humans and *Pan troglodytes*. Although the negative impact might be sharply reduced by the “ecological niche effect” mentioned above, using one clade-C1 genome to represent all clade-C genomes is destined to cause omissions. It has been observed that Clade D zooxanthellae also occupy a certain percentage of the *P. verrucosa* holobiont (**Video S1**), implying that a great deal of information is yet to be mined. Besides, the mixed *Symbiodinium* samples extracted from coral are still hard to isolate and culture in vitro currently. Acquiring genomes of specific *Symbiodinium* species precisely from coral-zooxanthellae holobionts can be described as the most essential challenge related to cross-kingdom regulation research.

In summary, our work integrates new full-length transcriptomes and small RNAs of four common and frequently dominant reef-building corals with a public *Symbiodinium C1* genome to explore coral-zooxanthella holobionts, shedding light on intracellular mutual regulation between these ancient heterotrophs and autotrophs. It reveals the miRNA-based anti-rejection mechanism used by zooxanthellae, proposes the concept of coral as a model organism for cross-kingdom regulation along with ecological niche effect, and offers a foundation for further cross-kingdom research.

We have added all our data, including gene annotation and miRNA information, to the CoralBioinfo database (http://coral.bmeonline.cn), in a gesture to build a high-quality multi-omics platform for further coral research. Multiple web services have been developed for it, most of which turn out to be more user-friendly to bioinformaticians familiar with “dry lab” skills. Data contribution requests for joint efforts are always welcome.

## Materials and Methods

### RESOURCE AVAILABILITY

#### Lead contact

Further information and requests for resources and reagents should be directed to and will be fulfilled by the lead contact, Chunpeng He, or by Zuhong Lu.

#### Materials availability

This study did not generate any new reagents.

#### Data availability

Raw data from our own full-length transcriptome sequencing and small RNA sequencing are available from NCBI (<u>PRJNA544778</u>).

Though we sincerely recommend acquiring data via our CoralBioinfo database (http://coral.bmeonline.cn) introduced in this paper, we have already uploaded necessary supplementary data to better serve researchers.

(https://seunic-my.sharepoint.cn/:f:/g/personal/230218818_seu_edu_cn/Eo4QciTXwQRBkYqcKEacEN8BRCKffoV-299v2HFILGF6lQ?e=dlQJx9)

### EXPERIMENTAL MODEL AND SUBJECT DETAILS

All coral samples were collected and processed in accordance with local laws for invertebrate protection and approved by the Ethics Committee of Institutional Animal Care and Use Committee of Nanjing Medical University (protocol code IACUC-1910003 and date of approval is 10 October 2019).

The species in the study were collected from the Xisha Islands in the South China Sea (latitude 15°40’–17°10’ north, longitude 111°–113° east).

The coral samples were cultured in our laboratory coral tank with conditions conforming to their habitat environment. All the species were raised in a RedSea^®^ tank (redsea575, Red Sea Aquatics Ltd) at 26°C and 1.025 salinity (Red Sea Aquatics Ltd). The physical conditions of the coral culture system are as follows: three coral lamps (AI^®^, Red Sea Aquatics Ltd), a protein skimmer (regal250s, Reef Octopus), a water chiller (tk1000, TECO Ltd), two wave devices (VorTechTM MP40, EcoTech Marine Ltd), and a calcium reactor (Calreact 200, Reef Octopus), etc.

## METHOD DETAILS

### RNA extraction

All RNA extraction procedures follow the manufacturers’ instructions. The total RNA was isolated with TRIzol LS Reagent (Thermo Fisher Scientific, 10296028) and treated with DNase I (Thermo Fisher Scientific, 18068015). The high quality mRNA was isolated with a FastTrack MAG Maxi mRNA Isolation Kit (Thermo Fisher Scientific, K1580-02). Samples were separated from healthy *Acropora muricata, Montipora foliosa, Montipora capricornis* and *Pocillopora verrucosa* to ensure that enough high quality RNA—more than 10 µg—could be obtained for a cDNA transcriptome library.

### Full-length transcriptome sequencing

Before establishing a cDNA transcriptome library, the quality of total RNA had to be tested. Agarose gel electrophoresis was used to analyze the degree of degradation of RNA and whether it was contaminated. A Nanodrop nucleic acid quantifier was used to detect the purity of RNA (OD260/280 ratio), a Qubit RNA assay was used to quantify the RNA concentration accurately, and an Agilent 2200 TapeStation was used to accurately detect the integrity of the RNA.

The Clontech SMARTer^®^ PCR cDNA Synthesis Kit (Clontech Laboratories, 634926) and the BluePippin Size Selection System protocol, as described by Pacific Biosciences (PN 100-092-800-03) were using to prepare the Iso-Seq library according to the Isoform Sequencing protocol (Iso-Seq).

The RNA-seq library was prepared according to the standard library construction for the Illumina HiSeq X Ten, using three biological replicates for each species of coral.

We used the PacBio Sequel II platform with SMRT (single molecular real time) sequencing technology and SMRTlink v7.0 software (minLength 50; maxLength 15,000; minPasses 1) to process sequencing samples (20). After polymer read bases were performed, the subreads.bam files were obtained by removing the joint and the original offline data with lengths of under 50 bp. The CCSs (circular consensus sequences) were obtained by putting the subreads.bam file through the CCS algorithm, which is self-correcting for the single molecule multiple sequencing sequence. Consequently, the full-length-non-chimera (FLNC) and nFL (non-full-length, non-chimera) sequences were identified by detecting whether CCSs contained 5’-primer, 3’-primer and poly-A. FLNC sequences of the same transcript were clustered with a hierarchical n * log (n) algorithm to obtain consensus sequences. The corrected consensus reads were polished from consensus sequences (Arrow polishing) using LoRDEC (21) v0.7 software using the RNA-seq data sequenced with the Illumina HiSeq X Ten platform. Using CD-HIT (22) software (-c 0.95 -T 6 -G 0 -aL 0.00 -aS 0.99), all redundancies were removed in corrected consensus reads to acquire final full-length transcripts and unigenes for subsequent bioinformatics analyses.

### Small RNA sequencing

A total amount of 3 μg of RNA per sample was used as input material for the small RNA library. Sequencing libraries were generated using the NEBNext^®^ Multiplex Small RNA Library Prep Set for Illumina^®^ (NEB, USA) following the manufacturer’s recommendations, and index codes were added to attribute sequences to each sample. Briefly, the NEB 3’ SR Adapter was directly and specifically ligated to the 3’ end of miRNA, siRNA and piRNA. After the 3’ ligation reaction, the SR RT Primer was hybridized to the excess of 3’ SR Adapter that remained free after the 3’ ligation reaction, transforming the single-stranded DNA adapter into a double-stranded DNA molecule. This step is important to prevent adapter-dimer formation, and dsDNAs are also not substrates for ligation mediated by T4 RNA Ligase 1, so they do not ligate to the 5’ SR Adapter in the subsequent ligation step. Next, the 5’ ends adapter was ligated to the 5’ ends of miRNAs, siRNA and piRNA. Then the first strand of cDNA was synthesized using M-MuLV Reverse Transcriptase (RNase H–). PCR amplification was performed using LongAmp Taq 2X Master Mix, SR Primer for Illumina and index (X) primer. PCR products were purified on an 8% polyacrylamide gel (100V, 80 min). DNA fragments corresponding to 140∼160 bp (the length of small noncoding RNA plus the 3’ and 5’ adapters) were recovered and dissolved in 8 μL elution buffer. Finally, library quality was assessed with the Agilent Bioanalyzer 2100 system using DNA High Sensitivity Chips.

The clustering of the index-coded samples was performed on a cBot Cluster Generation System using a TruSeq SR Cluster Kit v3-cBot-HS (Illumia) according to the manufacturer’s instructions. After cluster generation, the library preparations were sequenced on an Illumina HiSeq X Ten platform and 50bp single-end reads were generated.

Clean data for downstream analysis were obtained by removing reads with low quality, poly-N|A|T|G|C, and 5’ adapter contaminants. Reads without the 3’ adapter or the insert tag were cleaned as well.

### Gene function annotation

Gene function was annotated using the following databases: Nr (NCBI non-redundant protein sequences) (23), Nt (NCBI non-redundant nucleotide sequences), Pfam (Protein family) (24), KOG (Clusters of Orthologous Groups of proteins) (25), Swiss-Prot (A manually annotated and reviewed protein sequence database) (26), GO (Gene Ontology) (27) and KEGG (Kyoto Encyclopedia of Genes and Genomes) (14). We use BLAST 2.7.1+ in NCBI setting the e-value ‘1e-5’ in Nt database analysis, Diamond v0.8.36 BLASTX software, setting the e-value to ‘1e-5’ in Nr, KOG, Swiss-Prot and KEGG databases analysis, and HMMER 3.1 package for Pfam database analysis.

### Gene expression analysis

We used RSEM (28) to align Illumina reads to full-length transcripts and to quantify gene expression. Transcript names were converted to Nr IDs to build the multi-coral expression profile. EBSeq (29) was finally called by RSEM to identify differential expression patterns.

### miRNA identification

After quality control and length filtering (18–35 nt) of raw small RNA sequences, known miRNA alignment and novel miRNA prediction were conducted. For known miRNA alignment, miRBase was used as the reference, while mirdeep2 (30) and srna-tools-cli (http://srna-workbench.cmp.uea.ac.uk) were employed to identify miRNAs and draw their secondary structures. For novel miRNA prediction, small RNA tags were mapped to RepeatMasker (31) and Rfam (32) to remove tags originating from protein-coding genes, repeat sequences, rRNA, tRNA, snRNA, and snoRNA. Next, miREvo (33) and mirdeep2 were integrated to predict novel miRNA among the remaining unannotated tags. All miRNA targeting sites on zooxanthella genes were predicted with psRobot (34) (no mismatches allowed).

miRDP2 (35) was employed to identify *Symbiodinium* miRNAs in each coral separately after building the bowtie index of the *Symbiodinium C1* genome (36) downloaded from Reefgenomics (37). Initial outputs were corrected by genome annotation in which sequences with loci hitting CDS were removed. Then mirdeep2 using filtered sequences as inputs was run on all raw data to quantify miRNA expression as well to eliminate false negatives. All miRNA targeting sites were finally predicted with miRanda (38) (-sc 145 -en -10 -scale 4 -strict).

### Enrichment analysis

For each coral, the top 500 most highly expressed genes (that each entered the top 500 among three biological replicates more than once), and the zooxanthellae genes annotated by Nr and predicted target genes were all selected for GO/KEGG enrichment using clusterProfiler (39) (pvalueCutoff = 0.05, pAdjustMethod = ‘BH’, qvalueCutoff = 0.2).

### Website development

CoralBioinfo (http://coral.bmeonline.cn) was developed under the LEMP (Linux, Nginx, MySQL and PHP) stack. Web functions were implemented with PHP 7.2, while Nginx works as the web server. Most services have been built into Docker images, which are beneficial for cross-platform deployment and extended development.

All data are stored in MySQL tables with indexes to improve query efficiency. In addition, sequence data have been formatted into NCBI-BLAST local databases for alignment and download services.

## ADDITIONAL RESOURCES

Detailed database structure manuals along with user tutorials are all available at http://coral.bmeonline.cn/help.php.

API documents are available at http://coral.bmeonline.cn/api/.

## Supplemental information

**Fig S1. Annotation of four full-length coral transcriptomes with Nr database**. Horizontal axis is species (txid) and vertical axis is unigene number. The top three species with the highest number of annotated unigenes are *A. digitifera, E. pallida* and *N. vectensis*, which account for 92.8% in *A. muricata*, 89.8% in *M. foliosa*, 90.5% in *M. capricornis* and 85.9% in *P. verrucosa*.

**Fig S2. Supplemental results of gene expression analysis**. (a) Differential zooxanthella gene expression patterns among the four coral species studied. The horizontal axis is the pattern name and the vertical axis is the gene number. The top three patterns in quantity are Pattern05, Pattern02 and Pattern08, which represent differentially expressed genes in *M. foliosa, P. verrucosa* and *A. muricata*, while all Pattern01 genes identified as being co-expressed among species fail to meet the statistical significance criteria (PPDE>0.95). (b) Venn graph of top 500 most highly expressed genes among four species.

**File S1. Supplementary processing details of full-length transcriptomes**.

**File S2. Supplementary processing details of small RNAs**.

**Table S1. Complete results of miRNA-target gene enrichment analysis**. GeneIDs are CSG/T IDs in CoralSym (http://coral.bmeonline.cn/sym/).

**Table S2. Expression profiles of *Symbiodinium* miRNAs**. AM_x, MF_x, MC_x, PV_x are short for the biological replicates of *A. muricata, M. foliosa, M. capricornis*, and *P. verrucosa*. The expression level is quantified in RPM.

**Video S1. Demo of searching CoralTrait++ via CoralAPI**. CoralTrait++ is a manually-built simplified version of the Coral Trait database. It consists of 12 quantitative traits and six descriptive traits with all units standardized, and more than 45,000 records. It was revealed in this demo that clade D zooxanthellae also hold a certain percentage of the traits seen in the *Pocillopora verrucosa* holobiont.

**Data S1. Unigene sequences of *A. muricata, M. foliosa, M. capricornis*, and *P. verrucosa***. BLAST service is available at http://coral.bmeonline.cn/blast_unigene/.

**Data S2. Gene annotation of four corals from Nr, Nt, Pfam, KOG, Swiss-Prot, GO and KEGG databases**. All gene information is searchable at http://coral.bmeonline.cn/gene/.

**Data S3. CoralSym dataset including information of 11 novel *Symbiodinium* miRNAs and their targeting sites**. The format manual is available at http://coral.bmeonline.cn/sym/help.php #database-content. Exact searching of targeting information is accessible at /sym/search.php.

## Acknowledgments

We are grateful for Dr. Xiaojun Xia’s technical support in our production environment.

## Author Contributions

YZ: experiment, database, writing and editing. TH: data uploading. ZL & JC: reviewing. CH: supervision. XL: project approval. All authors contributed to the article and approved the submitted version.

